## Supplementary text 1 for "Absolute quantitation of serum antibody reactivity using the Richards growth model for antigen microspot titration"

Recently Pfeil and Herbály [5] proposed a linear model for polyclonal antibody-antigen reactions, which describes the initial phase of the reaction but does not reproduce the sigmoidal dependence of the fluorescent signal intensity related to the equilibrium concentration of the composed complex on the logarithm printed antigen concentration [1]. Here we modify the polyclonal linear model taking into account that the bound antibodies inhibit free antibodies to form a complex with nearby antigens [2], [3]. As a result of this steric hindrance, the equilibrium concentration of the complexes is reduced compared to that of the model of [5], and it makes the equilibrium concentration of the composed complex be a logistic function of the logarithm printed antigen concentration. As in the original model, we assume also in the modified model that both antibodies and antigens are monovalent. We also assume that any antigen is inhibited at most by one antibody.

Suppose that the serum contains $n$ different clones of antibodies with association reaction rate constants $k_{a_i}$, dissociation reaction rate constants $k_{d_i}$, equilibrium association constants $K_{A_i} = \frac{k_{a_i}}{k_{d_i}}$, and equilibrium dissociation constants $K_{D_i} = \frac{k_{d_i}}{k_{a_i}}$, and constant concentrations $[Ab_i]$ for $i = 1, \ldots, n$. Let $[Ag]$ denote the surface concentration of the printed antigen, and $[Ab_iAg]$ denote the surface concentration of complexes composed with the $i$th clone, and let $N_{Ab_iAg}$ the number of those complexes. Let a bound antibody inhibit nearby antigens of surface $S$, and let $S_0$ denote the surface of the antigen spot, and let $N_A$ be the Avogadro's number. Then the geometric probability of a free antigen to be inhibited is

$$P = \frac{S\left(N_{Ab_1Ag} + \ldots + N_{Ab_nAg}\right)}{S_0} = SN_A\left([Ab_1Ag] + \ldots + [Ab_nAg]\right).$$

The concentration of the available, i.e. free and not inhibited antigens is

$$\begin{aligned} [Ag]_{\mathrm{av}} &= [Ag] - ([Ab_1Ag] + \ldots + [Ab_nAg]) - P[Ag] \\ &= [Ag] - (1 + SN_A[Ag])\left([Ab_1Ag] + \ldots + [Ab_nAg]\right). \end{aligned} \tag{1}$$

We can modify the system of differential equations of [5] describing the kinetics of the polyclonal reaction by changing the free antigen concentration to the one of the available antigen to obtain system

$$\left.\begin{aligned} \tfrac{\mathrm{d}}{\mathrm{d}t}[Ab_1Ag] &= k_{a_1}[Ab_1][Ag]_{\mathrm{av}} - k_{d_1}[Ab_1Ag] \\ &\vdots \\ \tfrac{\mathrm{d}}{\mathrm{d}t}[Ab_nAg] &= k_{a_n}[Ab_n][Ag]_{\mathrm{av}} - k_{d_n}[Ab_nAg] \end{aligned}\right\},$$

$$\frac{\mathrm{d}}{\mathrm{d}t}\begin{pmatrix} [Ab_1Ag] \\ \vdots \\ [Ab_nAg] \end{pmatrix} = A\begin{pmatrix} [Ab_1Ag] \\ \vdots \\ [Ab_nAg] \end{pmatrix} + \begin{pmatrix} k_{a_1}[Ab_1] \\ \vdots \\ k_{a_n}[Ab_n] \end{pmatrix}[Ag], \tag{2}$$

where the matrix of the modified model is

$$A = \begin{pmatrix} -k_{a_1}[Ab_1]W - k_{d_1} & -k_{a_1}[Ab_1]W & \ldots & -k_{a_1}[Ab_1]W \\ -k_{a_2}[Ab_2]W & -k_{a_2}[Ab_2]W - k_{d_2} & \ldots & -k_{a_2}[Ab_2]W \\ \vdots & \vdots & \ddots & \vdots \\ -k_{a_n}[Ab_n]W & -k_{a_n}[Ab_n]W & \ldots & -k_{a_n}[Ab_n]W - k_{d_n} \end{pmatrix}$$

with $W = 1 + SN_A[Ag]$. The system is identical and the matrix has a form similar to those of the linear model for the polyclonal reaction, therefore applying Theorem 2 of [5] the equilibrium concentration of the $i$th complex is

$$[Ab_iAg]_{\mathrm{eq}} = \frac{[Ag]}{1 + (1 + SN_A[Ag])\sum\limits_{i=1}^{n} K_{A_i}[Ab_i]} K_{A_i}[Ab_i],$$

and that of the total complex is

$$[AbAg]_{\text{eq}} = \sum_{i=1}^{n} [Ab_iAg]_{\text{eq}} = \frac{[Ag] \sum\limits_{i=1}^{n} K_{A_i}[Ab_i]}{1 + (1 + SN_A[Ag]) \sum\limits_{i=1}^{n} K_{A_i}[Ab_i]}.$$

The equilibrium association constant of a polyclonal solution according to [4] is

$$K_A = \frac{1}{[Ab]} \sum_{i=1}^{n} K_{A_i}[Ab_i]$$

with the total antibody concentration $[Ab] = \sum\limits_{i=1}^{n} [Ab_i]$, thus

$$[AbAg]_{\text{eq}} = \frac{[Ag]K_A[Ab]}{1 + (1 + SN_A[Ag])\, K_A[Ab]} = \frac{[Ag][Ab]}{(K_D + [Ab]) + SN_A[Ag][Ab]}. \qquad (4)$$

This formula with $D = SN_A$ is formula (1) of the article. Rewriting formula (4) we obtain that the equilibrium concentration of the complex is a logistic function of the logarithm printed antigen concentration:

$$[AbAg]_{\text{eq}} = \frac{\frac{1}{SN_A}}{\frac{K_D + [Ab]}{SN_A[Ab]} \cdot \frac{1}{[Ag]} + 1} = \frac{\frac{1}{SN_A}}{1 + \frac{K_D + [Ab]}{SN_A[Ab]}\, e^{-\ln [Ag]}}. \qquad (5)$$

Finally, experience shows that the surface concentration of the printed antigen $[Ag]$ is proportional to its given volume concentration before printing $[Ag]_V$, i.e. $[Ag] = \alpha [Ag]_V$, and that the logarithm fluorescent intensity $\ln FI$ is proportional to the logarithm complex concentration $\ln[AbAg]_{\text{eq}}$, i.e. $\ln FI = \beta \ln[AbAg]_{\text{eq}}$, where $\alpha, \beta$ are positive constants. Hence we get

$$\begin{aligned} FI &= [AbAg]_{\text{eq}}^{\beta} = (SN_A)^{-\beta} \left(1 + \frac{K_D + [Ab]}{SN_A[Ab]}\, e^{-\ln[Ag]}\right)^{-\beta} \\ &= (SN_A)^{-\beta} \left(1 + \frac{1}{\beta}\, e^{-1 \cdot \left(\ln[Ag]_V - \ln \frac{\beta(K_D + [Ab])}{\alpha SN_A[Ab]}\right)}\right)^{-\beta}, \end{aligned}$$

which is a Richards function of $\ln[Ag]_V$ with parameters

$$A = (SN_A)^{-\beta}, \quad k = 1, \quad x_i = \ln \frac{\beta\,(K_D + [Ab])}{\alpha SN_A[Ab]}, \quad d = \frac{1}{\beta} + 1.$$
