## Supplementary material for "Absolute quantitation of serum antibody reactivity using the Richards growth model for antigen microspot titration": supplemetary text 2

**Supplementary text 2**

*Interpretation of the logarithmic form of the Richards curve as titration of chemical potential*

Chemical potential μ by definition is

$$\mu=\mu°+\ln(a) \qquad (S1)$$

where μ° is standard chemical potential, *a* is relative thermodynamic activity:

$$a=c\gamma \qquad (S2)$$

where *c* is molar concentration and γ is activity coefficient.

The chemical potential of a given antibody species is then described, after replacing *a* in (S1) with γ*c* from (S2)

$$\mu_{Ab} = \mu°_{Ab} + \ln\left(c_{Ab}\gamma_{Ab}\right) \qquad (S3)$$

$$\mu_{Ab} = \mu°_{Ab} + ln(c_{Ab}) + \ln\left(\gamma_{Ab}\right) \qquad (S4)$$

Omitting the standard term and raising both sides to the exponent, we obtain the proportionality

$$e^{\mu_{Ab}} \sim e^{\ln(c_{Ab})+\ln(\gamma_{Ab})} \qquad (S5)$$

We assume that the fluorescence intensity signals are a measure of the absolute thermodynamic activity of antibodies, $\lambda_{Ab}$

$$FI_{Ab} \sim \lambda_{Ab} = e^{\mu_{Ab}} \qquad (S6)$$

Therefore, we suggest that in the logarithmic form of Richards function with k=1

$$\ln\left(FI_{Ab}\right) = \ln(A) + \ln\left(\left(1+(d-1)e^{-(x-x_i)}\right)^{\frac{1}{1-d}}\right) \qquad (S7)$$

ln(A) is the logarithm of fluorescent intensity corresponding to the detection antibody with molar concentration $ln(c_{Ab})$ in S5, and the rest of the equation corresponds to logarithm of activity coefficient of antibody $\ln(\gamma_{Ab})$. However, we also propose that this second part of the equation also comprises activity coefficients and molar concentrations.

Distributing the negative sign in the exponent in S7, we get

$$\ln\left(FI_{Ab}\right) = \ln(A) + \ln\left(\left(1+(d-1)e^{x_i-x}\right)^{\frac{1}{1-d}}\right) \qquad (S8)$$

then assuming $1/(d-1)=\gamma_{Ag}^{\infty}$ (S9)

where $\gamma_{Ag}^{\infty}$ is constant for a given interaction at infinite Ab dilution, we obtain

$$\ln\left(FI_{Ab}\right) = \ln(A) + \ln\left(\left(1+\frac{1}{\gamma_{Ag}^{\infty}}e^{x_i-x}\right)^{-\gamma_{Ag}^{\infty}}\right) \qquad (S10)$$

By substituting $e^{x}=[Ag]$ and $e^{x_i}=[Ag]_i$

$$\ln\left(FI_{Ab}\right) = \ln(A) + \ln\left(\left(1+\frac{1}{\gamma_{Ag}^{\infty}}\frac{[Ag]_i}{[Ag]}\right)^{-\gamma_{Ag}^{\infty}}\right) \qquad (S11)$$

Comparing S4 and S11, the proposed activity coefficient of the measured antibody (highlighted in yellow) changes as a function of Ag concentration at inflection point $[Ag]_i$, concentration of antigen [Ag], and antigen activity coefficient $\gamma_{Ag}^{\infty}$ at infinite antibody dilution.

Overall, chemical potentials of the interaction partners are interdependent and are determined by the affinity distribution of antibodies in our measurement system.
