## supplementary figure S1 for "Absolute quantitation of serum antibody reactivity using the Richards growth model for antigen microspot titration"

Figure S1. Application of two-dimensional titration for the comparative measurement of anti-citrullinated antibodies. Four serum samples positive for anti-citrullinated IgG and IgA were tested for reactivity to VCP2 peptide [1]. Using the values of the parameters obtained by curve fitting, normalized, quantitative results are comparable not only for serum samples (A) but also for distinct antibody classes (B). Bar charts show mean $x_i$ and d values, and 95% confidence intervals.

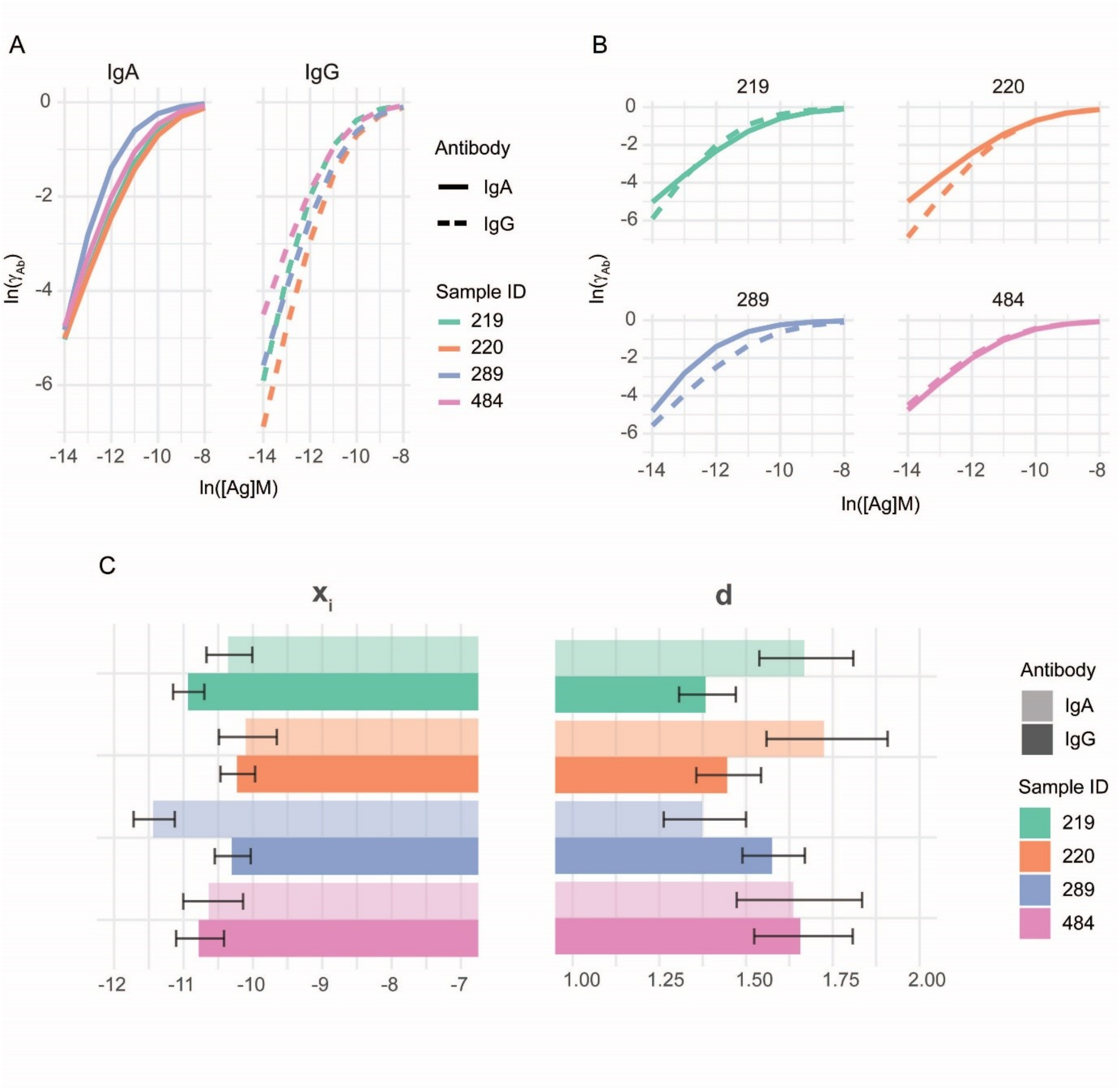
